# Symptoms expression of bakanae disease following seed treatment with phytohormone and metabolites

**DOI:** 10.1101/725978

**Authors:** Shireen A. Jahan Quazi, Sariah Meon, Zainal Abidin B.M. Ahmad, Hawa Jaafar

## Abstract

Gibberellic acid (GA_3_) phytohormone responsible for bakanae disease development is well-known and identified. But a number of secondary metabolites along with GA_3_, produced by the causal pathogen in relation to bakanae symptoms expression were unknown. Therefore, the aims of this research were to evaluate the symptoms expression analysis of bakanae disease by pre-seed treatment with pure (synthetic) phytohormones and metabolites in susceptible rice variety MR 211. The typical bakanae symptoms were evaluated by applying pure GA_3_, FA and MON either singly or in mixtures. It was confirmed that higher concentration of GA_3_ singly or with higher concentration of GA_3_ and MON in mixtures, caused unusual elongation of internodes. Plants became stunted when high concentration of FA was applied. Browning of leaves and stems, crown rot, root necrosis occurred and root length was decreased when mixtures of higher concentration of FA *MON*GA_3_ were used as pre-treatment. Similar observations were noted in plants inoculated with *F. proliferatum* at different score levels. The mechanisms of bakanae disease development through different symptoms expression in susceptible variety infected with *F. proliferatum* were identified.

## Introduction

*Fusarium fujikuroi*, causal agent of bakanae disease is known to produce gibberellic acid (GA_3_), fumonisin (FB1), monilifornin (MON), fusaric acid (FA) and beauvericin (BEA) in diseased plant [1-4]. GA_3_ is a growth promoting phytohormone but abnormal production of GA_3_ has been identified as a break-through in disease development and disease resistance or susceptibility in plants [5]. In addition, GA_3_ has been identified as being responsible for increase in plant height, whereas FA is responsible for decrease in plant height. Fungal metabolites MON, FB1 and BEA have been identified for causing phytotoxicity in plants rather than establishment of pathogenicity or disease symptoms expression in bakanae diseased plants [1, 6-7]. Besides *Fusarium fujikuroi, Fusarium proliferatum* is also identified as a causal agent of bakanae disease [8] and GA_3,_ FB1, MON, FA were also isolated from bakanae diseased plants infected with *F. proliferatum* [9]. It was also assumed that GA_3,_ MON and FA had strong influence on bakanae symptoms expression as a significant amounts of GA_3,_ MON and FA were isolated from the infected susceptible variety (data not presented here). Therefore, this study was carried out to verify the role of GA_3_, MON and FA on bakanae symptoms expression, following exogenous pre-treatment on seeds of susceptible variety MR 211 instead of inoculation with *F. proliferatum* over time.

## Materials and methods

### Soil mixture

Sand 40%, clay 30% and peat (PEATGRO) 30% were mixed well and sterilized at 120 °C for 90 min. Trays (28 cm × 21 cm × 6.5 cm) were filled with 2 kg of the sterilized soil mixture and used for sowing pre-germinated seeds.

### Chemicals used

Pure (Synthetic) GA_3,_ FA, and MON were used in this study. These chemicals were chosen based on positive relationship that were observed in relation to bakanae symptoms expression in susceptible variety MR 211 (data not presented here). Two concentrations of each chemical (GA_3_=15 µg/g, GA_3_=10 µg/g, FA= 400 µg/g, FA= 100 µg/g, MON= 120 ng/g, MON= 80 ng/g) were used in this study. The pure chemicals GA_3,_ FA, and MON were purchased from Sigma-Aldrich.

### Experimental layout and design

Pre-germinated seeds (soaked in water for 48 h) of susceptible variety MR 211 were soaked for 12 h in the probation hormone (GA_3_) and metabolite (FA and MON) solutions at two concentrations singly to find out the effective concentration suitable for causing infection or symptoms expression. This moderate pre-treatment period (12 h) was chosen in order to avoid the toxicity caused by the metabolites FA and MON treatment over prolonged periods at higher concentration used and to avoid death of plants at the very early stages of application ^6, 10, 11^. The two concentrations of each treatment were determined based on the highest and the lowest concentrations derived from bakanae diseased plants (data not presented here). The pre-treated seeds were sown in the sterilized soil in trays. Untreated pre-germinated seeds and pre-germinated seeds inoculated with *F. proliferatum* (10^6^ conidia/mL) used as controls were sown at the same time as phytohormone and metabolite pre-treated seeds. Seeds were sown in two rows per tray with 15 seeds per row. A total of 8 treatments along with 2 controls were used in this experiment. Each tray represented a single replication and trays were arranged in a completely randomized design with 3 replications per treatment (30 seeds per replication). All plants were maintained in a glasshouse with day and night temperatures of 30–35 °C and 23–30 °C, respectively, and watered daily. No fertilizer was applied to avoid any effects on the phytohormone and metabolites.

After the single pre-treatment trial, mixtures with two pre-treatment combinations and mixtures with three pre-treatment combinations were employed to observe the symptoms expression in mixtures with different treatment combinations. All procedures used were the same as in the single pre-treatment trial. Development of symptoms associated with the pre–treatments, either singly or in mixtures with two and three pre-treatment combinations was recorded weekly for 3 weeks.

### Sampling procedure and data collection

Plants were uprooted carefully without damaging the root tissues at the different sampling times. Sampling times were determined based on disease score levels as follows: level 1 (7 days after pre-treatment), level 3 (14 days after pre–treatment) and level 5 (21 days after pre-treatment). The stems, leaves (uppermost second leaf) and roots were separated, and stem and leaf lengths were measured using a measuring scale (1 m).

Roots were washed with water to remove soil and root length was recorded using the Root Scanner (Winrhizo V700, 2012b). Data expressed as percentage (%) increase or decrease in stem height and root length over the untreated control at each sampling time was analyzed (10 plants per replicate).

### Effects of pre-treatment with GA_3,_ FA, and MON on stem cell elongation (Histopathological study)

Random samples of stem sections from plants pre–treated singly with GA_3_ (15 mg/L), FA (400 mg/L), and MON (120 µg/L), in mixtures with GA_3,_ (15 mg/L) * FA (400 mg/L)* MON (120 µg/L), the untreated control and the diseased control were observed after 14 days of pre-treatment applied. Plants showing typical symptoms of bakanae disease in relation to disease score level 1 and 3 were collected and studied. Stems were cut into 1 cm sections and prepared for scanning electron microscope analysis. Average cell lengths in the longitudinal stem sections of the plants grown from pre-treated seeds and inoculated plants were compared with those in the untreated control plants. Average cell length was measured from 30 randomly selected cells from 3 sets of samples (10 cells/sample set) observed under SEM.

### Analysis of symptoms expression in plants pre-treated with pure GA_3,_ FA and MON applied singly or in mixtures

Symptoms expression were analysed in plants pre-treated with pure GA_3,_ FA, and MON at two concentrations applied singly and in mixtures with two and three treatment combinations at stipulated disease scoring levels (score 1, 3 and 5) on percent increase or decrease basis in comparison with diseased control plants. Data were analysed using SAS software (Version 9.2).

## Results

### Effects of GA_3_, FA and MON applied singly

Both GA_3_ concentrations increased stem height, whereas the other pre-treatments decreased stem height after 7 days of pre-treatment. Highest stem height decrease was observed in pre-treatment FA (400 mg/L) after 7 days of pre-treatment. It was also observed that the increasing trend in stem height in GA_3_ pre-treatment declined after 7 days to after 21 days. In contrast, pre-treatment with MON (120 µg/L) showed increasing stem height after 14 days of pre–treatment and was found to continue increasing after 21 days which was similar to pre– treatment with GA_3_ (15 mg/L). Percent decrease in stem height declined in pre–treatment with FA (400 mg/L) over time. Stem height increase/decrease following the treatment used were observed in Figure 1.

**Figure 1.**
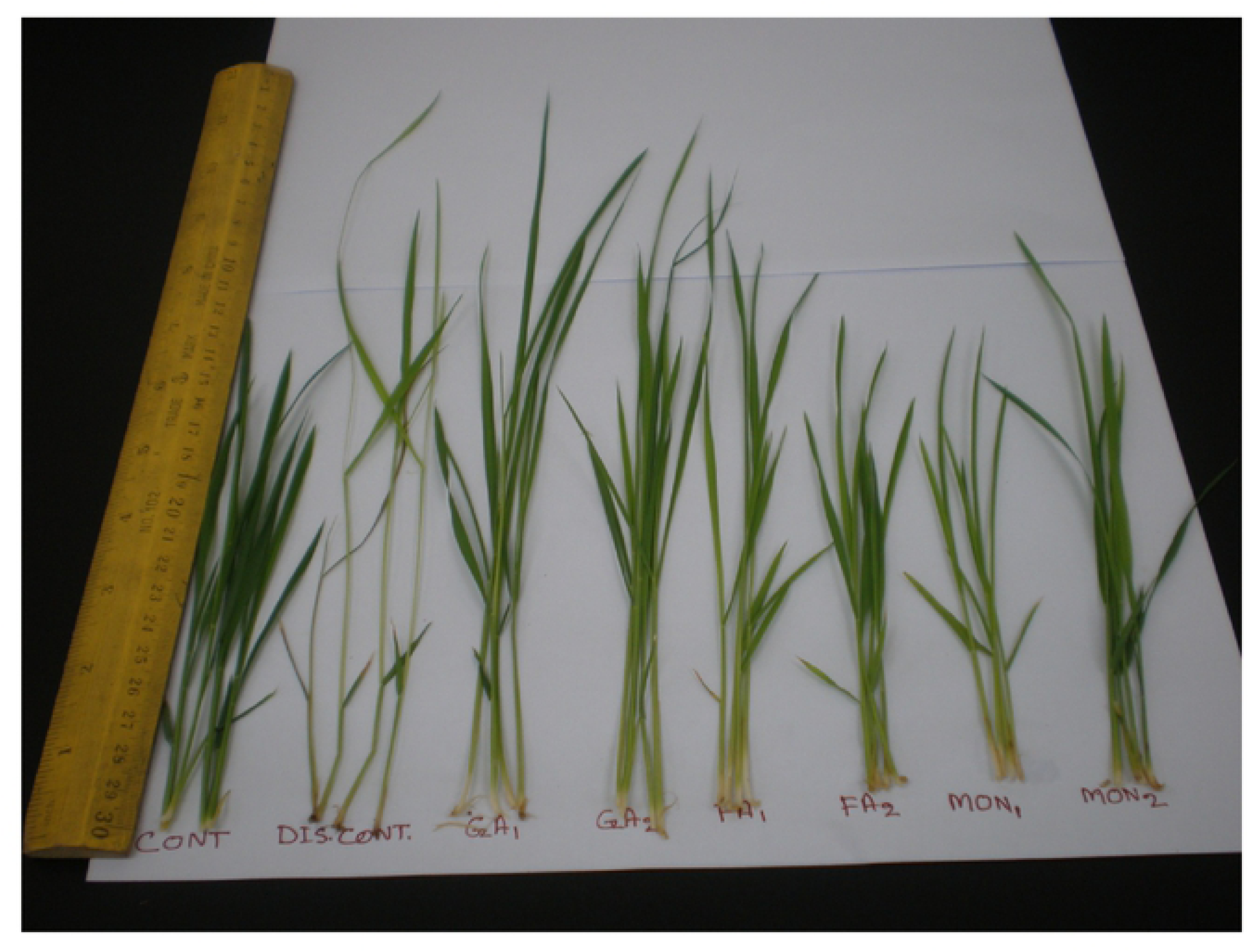
Stem height increase or decrease in plants following pre-seed treatment with GA_3_, FA, and MON singly in comparison with disease free (control) and disease control (inoculated) plants after 14 days.

These results were further confirmed by the histopathological study comparing cell lengths elongation with the different treatments. Stem cell length (average) was elongated (61.33 µm) in plants pre-treated with higher concentration of GA_3_ and was comparable to average stem cell length in diseased control plants (63.47 µm) after 14 days of pre-treatment and inoculation, respectively (Figure 2 a and b). Average cell length of stems was elongated somewhat (47.6 µm) when pre-treated with higher concentration of MON compared to average stem cell length of healthy control plants (45.33 µm) after 14 days (Figure 2 c and d).

**Figure 2:**
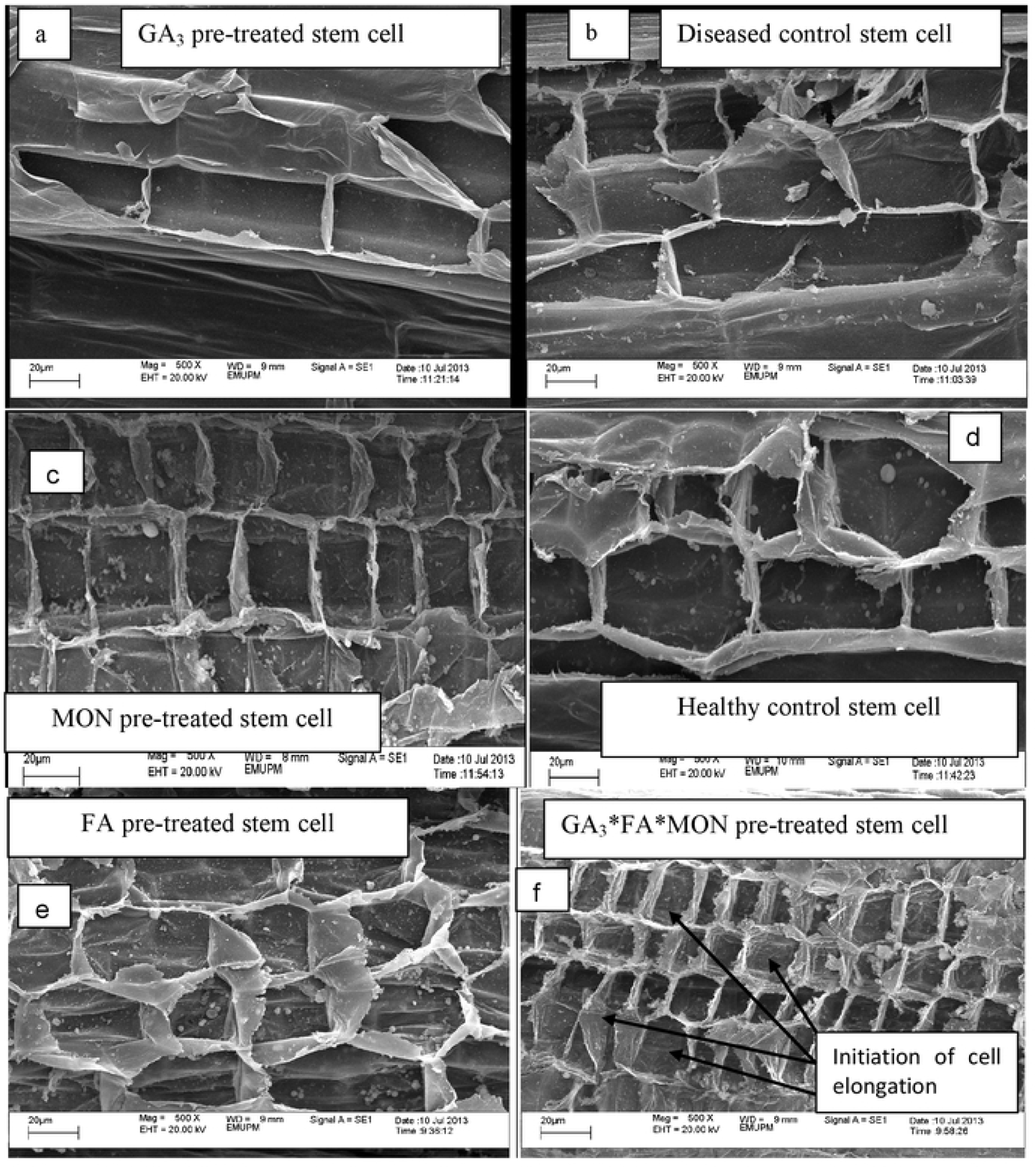
SEM micrograph of stem cell length in GA_3_ (15 mg/L), FA (400 mg/L) and MON (120 µg/L) pre-treated plants singly or in combination in comparison with diseased and healthy control plants. [GA_3_ pre-treated stem cells (a), Stem cells of diseased control plants (b), MON pre-treated stem cell (c), Stem cells of healthy control plants (d), FA pre-treated stem cell (e) and Stem cells in mixtures of pre-treatment with GA_3_ (15 mg/L)* FA (400 mg/L)* MON (120 µg/L) (f)]

In contrast, pre-treatment with FA showed stunting in stem height, which may be due to the shorter stem cell length. This percentage decrease in stem height observed was probably due to the higher cell division that occurred in plants at disease score 1, which caused shorter cell length (40 µm) in stems as compared to the stem cell length of healthy control plants (45.33 µm) observed in the histopathological study (Figure 2 e and d). From these result, it was evident that higher concentrations of GA_3_ and FA were responsible for the stem height increase and decrease, respectively, whereas the higher concentration of MON was responsible for marginal stem height increase after 14 days and onwards.

### Effects of GA_3_, FA and MON applied in combinations

In mixtures with two treatment combinations, seeds pre-treated with GA_3_ (15 mg/L)* MON (120 µg/L) resulted in the highest (%) increase in stem height after 7 −14 days and was found to the next to the diseased control plants. All combinations with GA_3_ and MON increased stem height after 14 days. Therefore, it was apparent that both GA_3_ and MON had a synergistic effect on stem height increase. Thus, at the disease score level 3, stem height was increased after 14 days, and this increasing trend was observed in pre-treatment combination treatments with GA_3_ (15 mg/L)* MON (120 µg/L). In contrast, there might be an antagonistic response between FA and GA_3_ and stem height increase or decrease was observed in relation to higher concentration of involvement with GA_3_ or FA vice versa. Stem height increase in plants of pre-treated seeds with GA_3_ (10 mg/L)* FA (400 mg/L) and in inoculated plants (diseased control) were observed to be almost similar after 7days. Thus, stunting observed in plants after 7 days of inoculation might be due to the effect of higher concentration of FA present in plants at the disease score level 1 (after 7 days of inoculation).

The mixtures with three pre-treatment combinations of FA (400 mg/L)* MON (80 µg/L)* GA_3_ (15 mg/L) and FA (100 mg/L)* MON (120 µg/L)* GA_3_ (15 mg/L) increased stem height after 7 days. This is largely due to the effect of GA_3_ (15 mg/L) singly as the treatment increased stem height early at 7 days after pre–treatment. Although stem cell length was observed to be shortened in this pre–treatment compared to control plants (Figure 2 f and d) but stem height increase after 14 days in this pre–treatment was attributed to the effect of excessive cell divisions in this pre-treatment.

Along with increases in stem height, leaf browning, stem browning, crown rot and root necrosis were observed in plants pre-treated with MON at both concentrations after 21 days. Browning was found to be more prominent at the lower concentration of MON (80 µg/L) applied singly as pre–treatment, whereas crown rot and root necrosis were more prominent at the higher concentration of MON (120 µg/L) (**Figure 3**). The browning symptoms observed in leaves and stems were similar to the symptoms on plants infected with *F. proliferatum* (diseased control), and could easily be distinguished from the symptoms on untreated control (healthy) plants (**Figure 3**).

**Figure 3:**
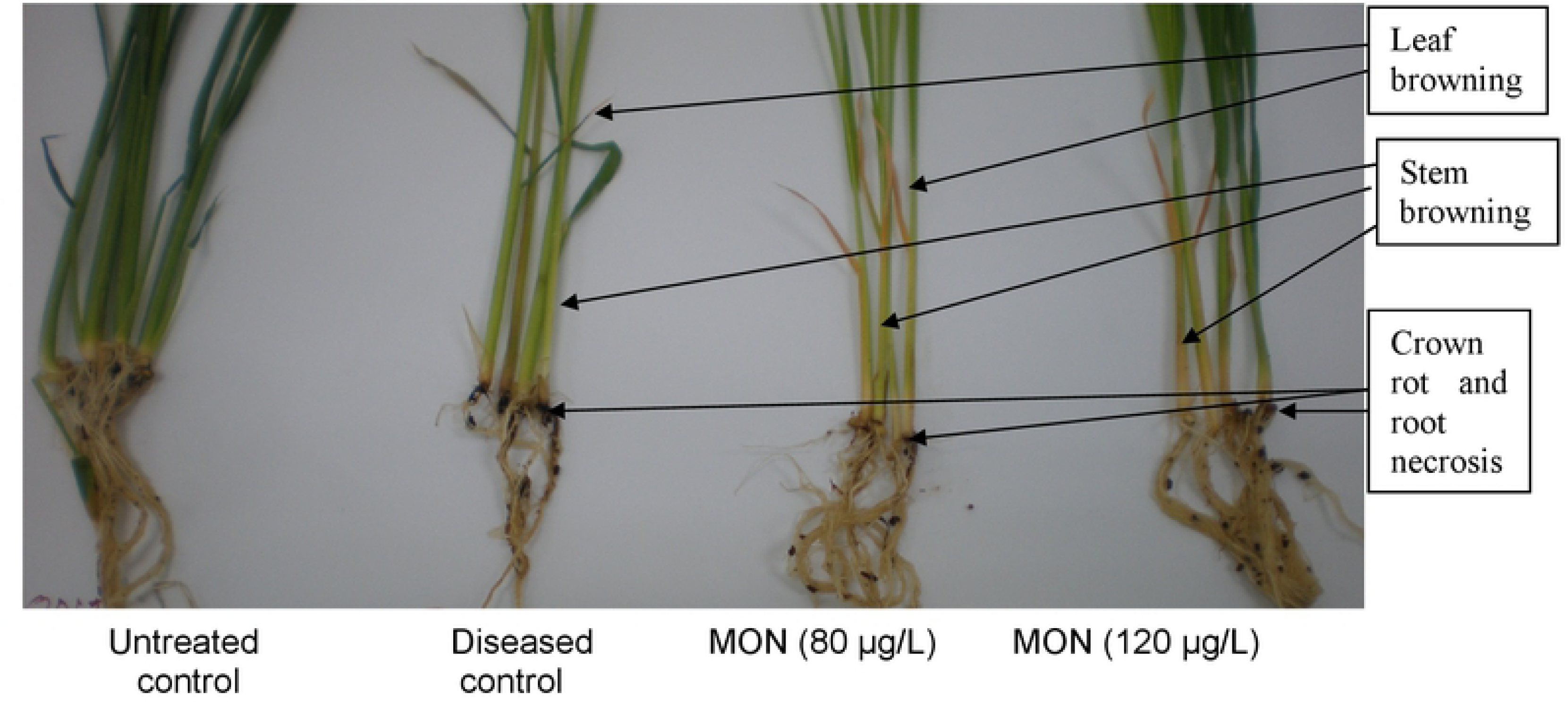
Plants pre-treated with MON singly and diseased control plants showing leaf browning, stem browning, crown rot and root necrosis in comparison with untreated (healthy) control plants after 21 days.

The highest decrease in root length was observed after 7 days, while it was found to increase after 14 days in all pre-treatments along with the diseased control. The highest decrease in root length was observed in pre–treatment with GA_3_ (15 mg/L) followed by the diseased control and GA_3_ after 7 days. Both concentrations of MON increased root length after 21 days of pre-treatment but with necrotic lesions.

In mixtures with two pre-treatment combinations the highest root length increase was observed in mixtures with GA_3_ (15 mg/L)* MON (120 µg/L) after 14 days (Figure 4a). It was also observed that both concentrations of MON caused crown rot and root necrosis in mixtures with GA_3_ in plants pre-treated with two pre-treatment combinations (Figure 5 a). It was also apparent that MON in mixtures either with GA_3_ or with FA caused prominent crown rot and root necrosis compared to that in mixtures with FA and GA_3_ (Figure 4 b and c).

**Figure 4:**
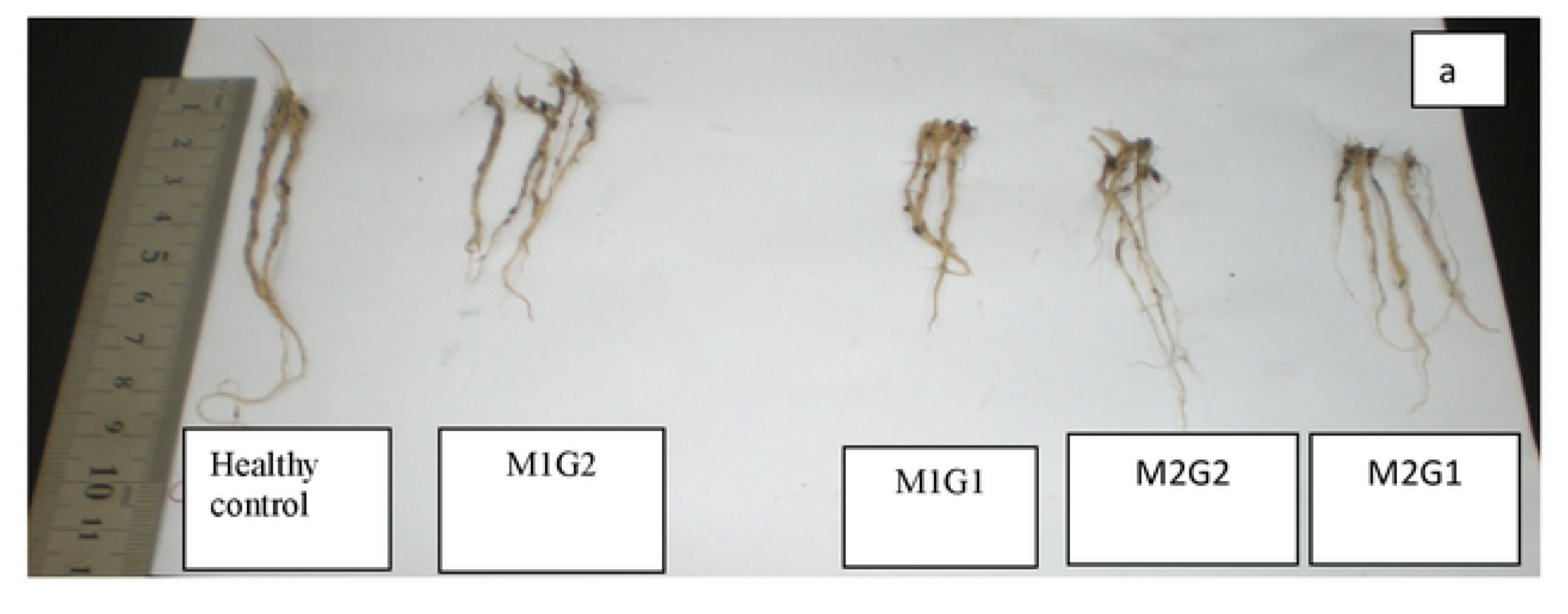
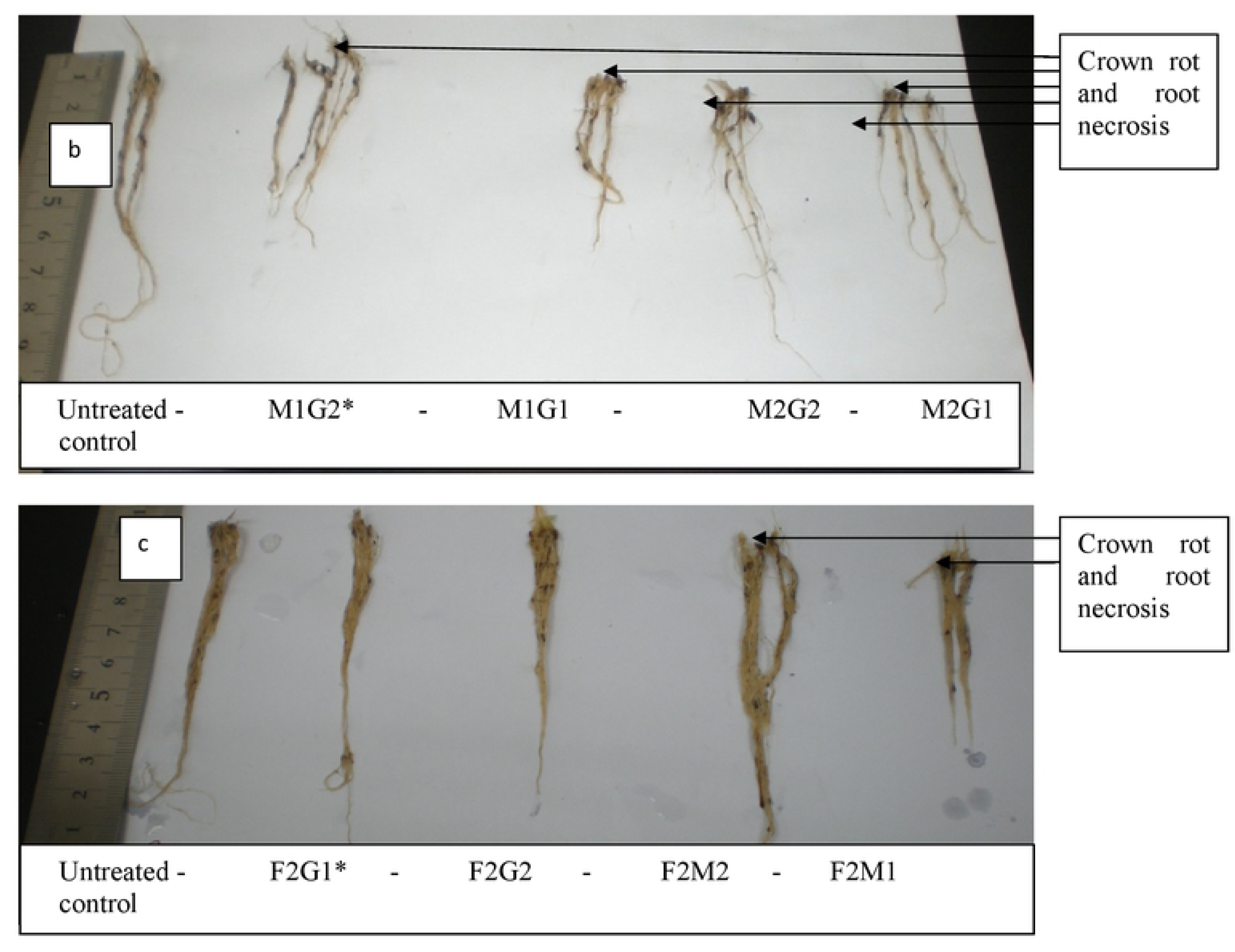
Plant roots showing different architectures when seeds were pre-treated with two treatment mixtures. [mixtures with MON and GA_3_ (a) and mixtures with FA and GA_3_ and mixtures with FA and MON (b)] *G1= GA_3_ (10 mg/L), G2= GA_3_ (15 mg/L), F1= FA (100 mg/L), F2= FA (400 mg/L), M1= MON (80 µg/L), M2= MON (120 µg/L)]. *MON = moniliformtn, GA_3_ = gibberellic acid, FA= fusaric acid

Crown rot and root necrosis was also observed to be severe in mixtures with three pre-treatment combinations in **GA**_**3**_ (15 mg/L)* MON (120 µg/L)* FA (400 mg/L) (Figure 5).

**Figure 5.**
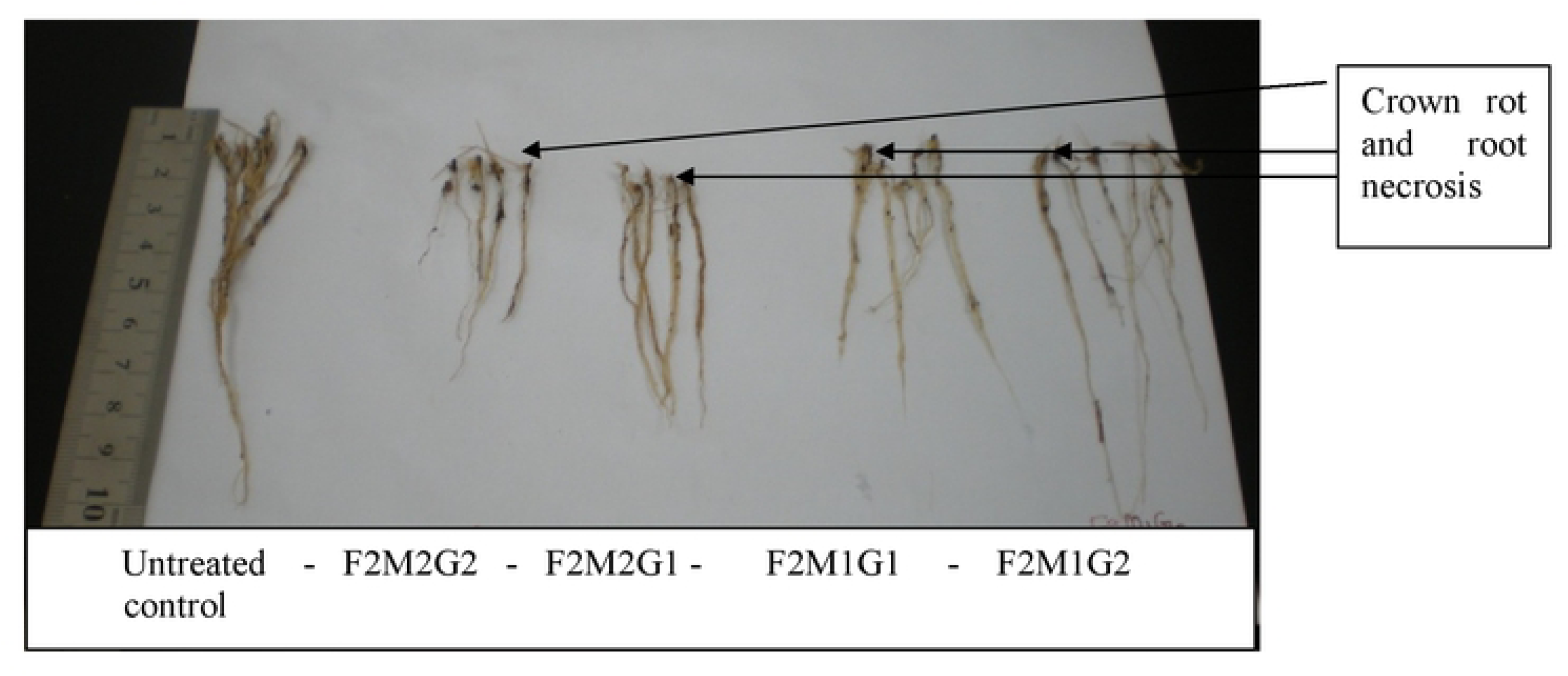
Crown rot and root necrosis due to the effect of MON in mixtures with three pre-treatment combinations. {*G1= GA_3_ (10 mg/L), G2= GA_3_ (15 mg/L), F1= FA (100 mg/L), F2= FA (400 mg/L), M1= MON (80 µg/L), M2= MON (120 µg/L)}. *MON = moniliformtn, GA_3_ = gibberellic acid, FA = fusaric acid

From the scenario in pre–treatments, either singly or in mixtures, it was observed that higher concentrations of GA_3_ and MON had a direct influence on increase in stem height. In contrast, FA singly in any concentration or at higher concentrations in mixtures with GA_3_ and MON decreased stem height.

## Discussion

Stem height increase in bakanae diseased plants at the disease score level of 3 (14 days after inoculation) was found to be dependent on higher amount of GA_3_ singly or in mixtures with MON. Although stem height increase (%) was observed to be lower in pre-treatment GA_3_ (15 mg/L)* MON (120 µg/L) compared to diseased control plants after 14 days but it was presumed to be due to slow down of GA_3_ activity with time. Moreover, increase in GA_3_ concentration was initiated in the diseased control plants after 7 days of pathogen inoculation and reached a maximum concentration in infected plants after 14 days, whereas pre-treatment with GA_3_ responded with a maximum level after 7 days. Similar observations were reported by other researchers, where GA_3_ concentration was increased to its highest level in plants after 1 week of application and subsequently decreased with time [12].

In contrast, pre-treatment with higher concentrations of FA singly or in mixtures with MON decreased stem height after 7 days and was reflected in bakanae diseased plants where stunting was observed at the disease score level of 1 after 7 days of inoculation. At the disease score level of 5, stem heights were found to decrease to some extent probably due to the effect of GA_3._ A possible explanation for this was that the increasing rate of GA_3_ was slowed down and FA was increased significantly due to antagonistic effect with each other with time in infected plants as observed after 21 days of pre–treatment. This is the first report that complex bakanae symptoms associated with the combination effect of increased GA_3_ levels along with metabolites FA and MON produced by the pathogen in infected plants, rather than solely dependent on increased levels of GA_3_ in infected plants. bakanae symptoms expression due to phytohormonal and metabolites effect is as illustrated in the Figure 6.

**Figure 6:**
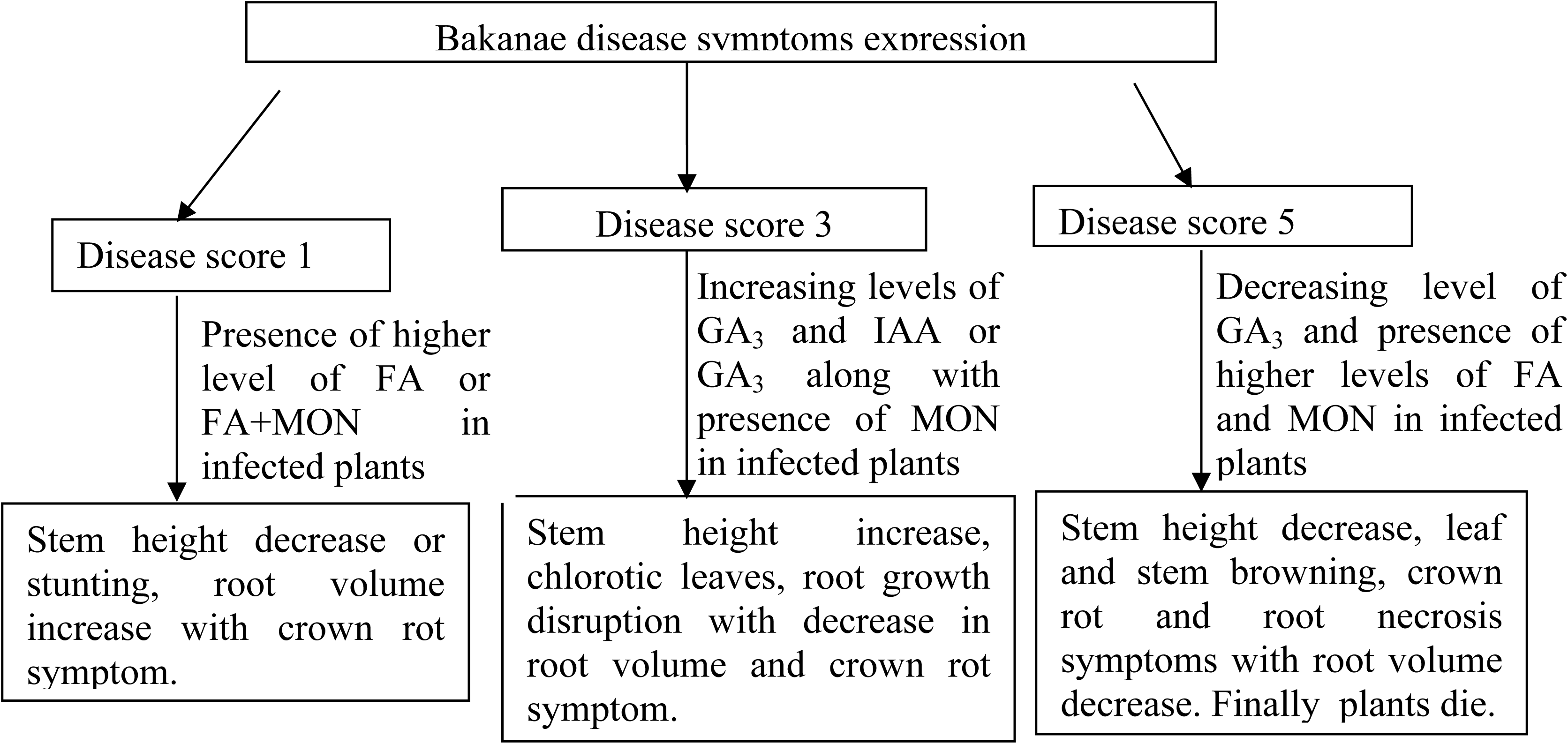
Flowchart of Bakanae symptoms expression due to phytohormonal and metabolites effect. (GA_3_= gibberellic acid, IAA= indole acetic acid, FA= fusaric acid, MON= moniliformin).

Stem height increase (%) was reliant on elongation of cell length as well as an increase in cell division inside the plant tissues as observed by several researchers. Other researchers reported that cell division was mainly influenced by GA_3_ application compared to cell elongation ^11^. The authors assumed that IAA might have an influence on plant height increase after cell division occurred with GA_3_ application. Later, other researchers explained that GA_3_ pre-treated plants stimulated IAA synthesis in plant cells first and IAA had an effect on cell elongation and this cell elongation was increased in aged tissues compared to young tissues [14-15].

Thus, the increase in stem height in bakanae diseased plants was due to the combined effect of cell division and cell length elongation. The mechanism is similar as observed in infected plants with *F. proliferatum* which induced cell division first due to increase levels of GA_3_ and thereby stimulated IAA production in infected plants. Thus, IAA in infected plants influenced cell enlargement in bakanae diseased plants. It was also established that IAA levels increased in bakanae diseased plants infected with *F. proliferatum* when GA_3_ was increased [9]. Several researchers have also supported that GA_3_ and IAA play their role synergistically in plants [15-17]. This higher level of GA_3_ along with increased IAA levels at the disease score level of 3, caused stem height increase compared to control plants after 14 days of inoculation. Although some stem height increase and cell enlargement was observed in MON pre-treated stem cells, but this response in relation to IAA increase has not been reported yet.

In the three pre-treatment combination FA (400 mg/L)* MON (120 µg/L)* GA_3_ (15 mg/L) stem height was increased (after 14 days) although shorter stem cell length (36.53 µm) was observed compared to the control (45.33 µm) in the histopathological study. The shorter cell length in this three pre-treatments mixture was probably due to an increase in cell division as a result of FA (400 mg/L) and/or GA_3_ (15 mg/L) and observed in the histopathological study after 14 days of inoculation. Moreover, it was reported that IAA was increased in plants at the second week after GA_3_ application ^10^, and that the increased IAA was reflected by a higher increase in stem height after 14 days with pre-treatment FA (400 mg/L)* MON (120 µg/L)* GA_3_ (15 mg/L). Again the shorter stem cell length (36.53 µm) was the average of 30 randomly selected stem cells pre-treated with FA (400 mg/L)* MON (120 µg/L)* GA_3_ (15 mg/L). Although some stem cells were initiated to elongate in this pre-treatment as observed in Figure 5.2e whereas, others were in initial stage of elongation and did not reflect individual cell enlargement in histopathological study where measurement was on average cell length. Additionally, stem height increase was slowed down after 7 days due to antagonistic effect of higher concentration of FA with GA_3_ and after 14 days stem height increased due to the effect of higher concentration of MON and GA_3_ combination.

In contrast, plant height stunting was found to be mainly due to the effect of FA rather than GA_3_. This observation is supported by other researchers as well. Other researcher reported that plant stunting or rossetting occurred due to low concentration of GA_3_ in plants ^16^. It was also observed that high levels of FA accumulation in plants resulted in stunted plant height and decreased root length in tomato [7, 19-20]. The lower amount of GA_3_ (8.9 µg/g fresh wt.) as determined in the disease score level of 1 resulted in stunting of plants after 7 days of pre-treatment. Although metabolic activity was not determined in this experiment but other researchers repoted that the stunting mechanism is mainly due to changes in metabolic activity in plants, including speeding up of the lipid peroxide activity, inhibition of ATP synthesis or decreased ATP levels [6, 19-20]. compared to control plants. Moreover, higher concentrations of FA and MON also resulted in plant height reduction as was observed in jimsonweed plants when pure FA and MON were applied ^7^. Thus, lower amount of GA_3_ and/or in mixtures with higher amounts of FA and MON contributed to the cessation and collapse of plants growth at the disease score level of 5 that was observed after 21 days of pre-treatment.

Crown rot and root necrosis due to higher concentration of MON in infected plants at later growth stages were observed. Similar symptoms have also been observed by other researchers in different plant species [7, 21]. Brown to pink discoloration of leaves at a disease score level of 5 may be due to higher concentration of MON present in plants infected by *Fusarium proliferatum.* It was also observed pink ear rot in maize when a higher concentration of MON was associated with the ear rot causal pathogen *Fusarium suubgltinans* [22]. In addition, earlier researcher reported that MON had toxic effects with growth reductions of coleoptiles as well as vein chlorosis and necrosis in corn and tobacco callus ^19^. MON was also found to cause cytoplasmic disruptions and abnormal mitosis that resulted in “a disruption of the spindle apparatus” in infected plant cells ^19, 21^. In contrast, reported that browning of stalk, leaf, and ear of rice was due to the effect of fumonisin B1 when inoculated with *F. proliferatum* ^22^. However, the authors did not isolate other mycotoxins produced by the fungus in the rice plants infected with *F. proliferatum* nor compared the symptoms treated with pure fumonisin B1. More recently provided evidence that FB1 had no pathogenicity effects on bakanae symptoms development ^1^. The browning or pinkish symptom associated with a disease score 5 was evaluated by applying pure MON to germinating seeds of susceptible variety MR 211, and similar symptoms were observed as in plants inoculated with *F. proliferatum.* Therefore, it was confirmed that browning and pinkish discoloration of leaves and stems as well as crown rot and root necrosis were associated with MON effect. Higher concentration of FA, MON and less GA_3_ accumulation in infected plants resulted in dead plants over time at a disease score level of 5, and this was attributed to inhibition of photosynthesis and toxic effects of MON and FA in plant cells.

